# Sub-second fluctuations in extracellular dopamine encode reward and punishment prediction errors in humans

**DOI:** 10.1101/2023.02.24.529709

**Authors:** L. Paul Sands, Angela Jiang, Brittany Liebenow, Emily DiMarco, Adrian W. Laxton, Stephen B. Tatter, P. Read Montague, Kenneth T. Kishida

## Abstract

In the mammalian brain, midbrain dopamine neuron activity is hypothesized to encode reward prediction errors that promote learning and guide behavior by causing rapid changes in dopamine levels in target brain regions. This hypothesis (and alternatives regarding dopamine’s role in punishment-learning) has limited direct evidence in humans. We report intracranial, sub-second measurements of dopamine release in human striatum measured while volunteers (i.e., patients undergoing deep brain stimulation (DBS) surgery) performed a probabilistic reward- and punishment-learning choice task designed to test whether dopamine release encodes only reward prediction errors or whether dopamine release may also encode adaptive punishment-learning signals. Results demonstrate that extracellular dopamine levels can encode both reward and punishment prediction errors, but may do so via by independent valence-specific pathways in the human brain.

**One-Sentence Summary:** Dopamine release encodes reward and punishment prediction errors via independent pathways in the human brain.

## Main Text

Dopamine neurons are critical for mammalian brain function and behavior (*1*), with changes in dopaminergic efficacy believed to underlie a wide range of human brain disorders including substance use disorders, depression, and Parkinson’s disease (*2–5*). The basic function of dopamine neurons is hypothesized to be to encode information about errors in an organism’s expectations about rewarding outcomes – so-called reward prediction errors (RPE; *6, 7*). Specifically, in non-human animal research, it has been shown that dopamine neurons encode “temporal difference” RPEs (TD-RPE; *6–13*), an optimal learning signal derived within computational reinforcement learning theory (*14*) and that has recently been central to major advances in the development of deep learning artificial neural networks capable of autonomously achieving human expert-level performance on a variety of tasks (*15–18*).

Decades of non-human animal research supports the idea that dopamine neurons encode RPEs in the mammalian brain (*6–13*; see *10* for review); however, in humans, direct evidence is limited. There is clear evidence in humans that changes in the firing rate of putative dopamine neurons encode RPEs (*19*), and regions rich in afferent dopaminergic input show changes in blood oxygen-level-dependent signals consistent with physiological processing of RPEs (*20–22*). Still, due to methodological limitations, these experiments cannot provide direct evidence that dopamine release in target regions encodes RPEs. In rodents, sub-second changes in extracellular dopamine levels in the striatum have been measured using fast scan cyclic voltammetry (FSCV) and rapid-acting, genetically encoded fluorescent dopamine sensors (e.g., dLight, GRAB; *23, 24*), revealing that dopamine levels reflect RPEs (*11–13*) but also respond to diverse affective stimuli (e.g., drug-predictive cues; *25, 26*) and vary with specific recording location (*27*) and task demands (e.g., effort costs; *28*). Consistent with this, rodent and non-human primate studies have shown that changes in dopamine neuron firing rate may also encode aversive prediction errors (*12, 29–33*). Relatedly, non-invasive human neuroimaging experiments suggest that reward and punishment prediction error signals are represented in dopamine-rich regions during learning about appetitive and aversive consequences (*34–37*).

Recently, studies leveraging the ability to directly measure dopamine release in the human brain with high temporal resolution have revealed that sub-second changes in dopamine levels reflect both actual and counterfactual error signals during risky decision-making (*38, 39*), the average value of reward following a sequence of decisions (*40*), and non-reinforced, though goal-directed, perceptual decision-making (*41*). In experiments where RPEs could be estimated (*38, 39*), dopamine levels seemed to entangle actual and counterfactual information (i.e., outcomes that “could have been” had a different choice been made) for both gains and losses, resulting in a superposed value prediction error signal (*38*). These results suggest the hypothesis that extracellular dopamine fluctuations encoding of reward and punishment prediction errors could be derived from independent streams of information processing, allowing these signals to be efficiently combined or differentiated by downstream neurons in the striatum (*42*).

We sought to determine whether dopamine release in human striatum specifically encodes TD-RPEs in humans as initially suggested by foundational work in non-human primates (*6,7*). We also sought to test an alternative hypothesis that dopamine release in these same loci also encodes punishment prediction errors, the possibility of which remains debated (*12, 29–33*). To test these hypotheses, we used human voltametric methods (*38–41, 43*) while participants performed a decision-making task (Fig. 1A) that allowed us to disentangle the impact of rewarding and punishing feedback on dopamine release and choice behavior. This approach allowed us to monitor dopamine release (Fig. 1B) while participants learned from rewarding as well as punishing feedback. The specific task design (*43*) allowed us to test two different reinforcement learning models that express the mutually exclusive hypotheses that dopamine release encodes reward- and punishment-prediction errors via 1) a unidimensional valence system, versus 2) a valence-partitioned system (*44*) whereby appetitive and aversive stimuli are processed by independent systems, thereby allowing learning of co-occurring though statistically independent appetitive and aversive stimuli (fig. S1; *43*).

**Figure 1.**
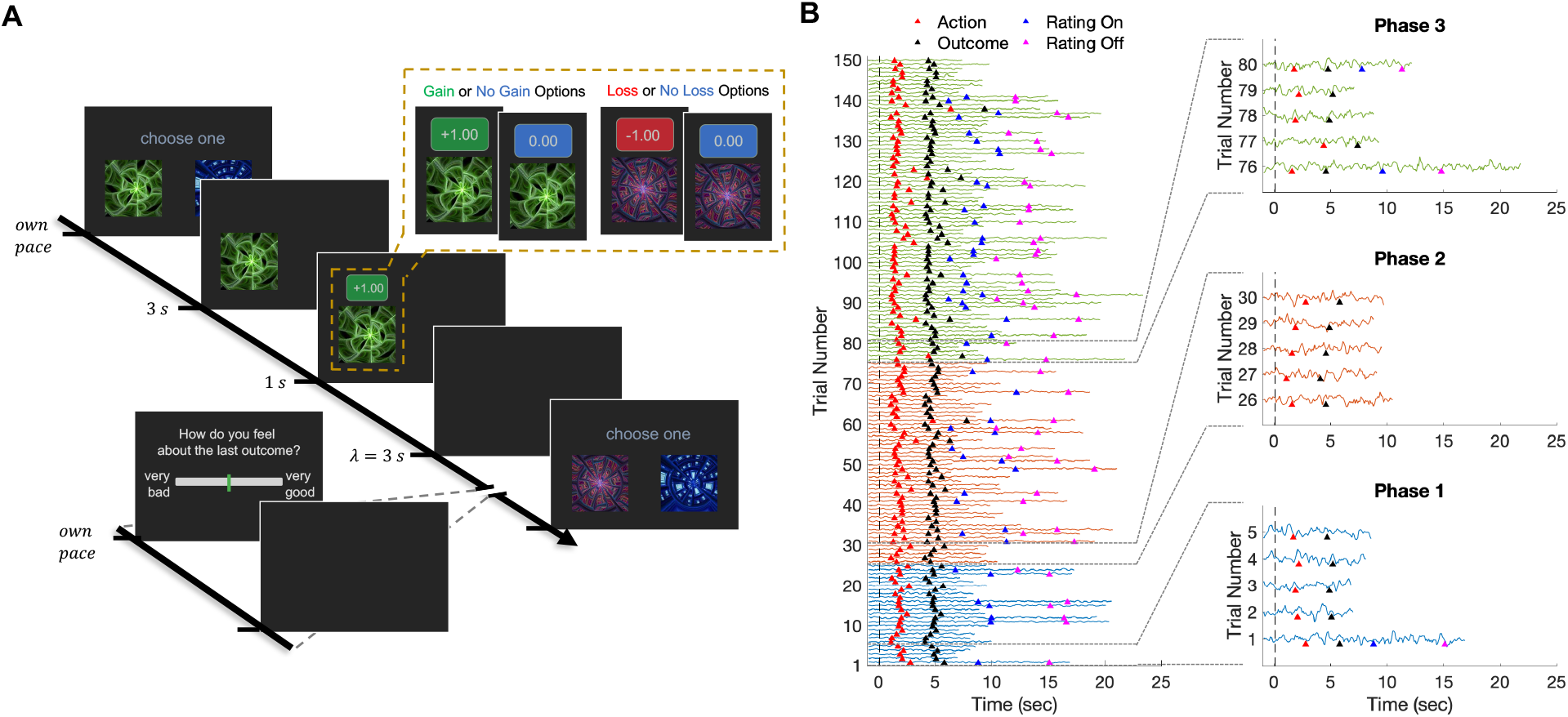
Probabilistic reward and punishment task and associated trial-by-trial dopamine time series recorded via human voltammetry. (**A**) Schematic of a trial from the choice task. **(B)** Trial-by-trial time series of caudate dopamine levels recorded from a single participant, with time series colored according to task phase; vertical dashed line indicates when the choice options were presented on each trial, and colored markers indicate trial events of interest.

### Human voltammetry experimental design

Participants (N=3) were adult patients diagnosed with essential tremor (ET) who consented to undergo deep brain stimulation (DBS) electrode implantation neurosurgery (*43*). Prior to the day of surgery, all participants provided written informed consent (*43*) to participate in the research procedure after deciding to undergo the clinical procedure. The neuroanatomical target of DBS lead implantation surgery for patients with ET is the ventralis intermediate nucleus of the thalamus, and this surgery includes micro-electrode recording within the caudate nucleus – a major site for dopaminergic innervation and dopamine release. Notably, the pathophysiology of ET is thought to not involve disruptions of the dopaminergic system (*45*). Prior to implanting the DBS lead, a carbon-fiber microelectrode is used for voltammetric recordings along the trajectory that the DBS lead may be placed (*38–41, 43*). In the present work, the carbon-fiber microelectrode was placed in the caudate, and dopamine measurements were sampled once every 100msec while participants performed the reward and punishment learning task. Following the research procedure, the carbon fiber microelectrode is removed and the DBS electrode implantation surgery is completed. Importantly, no change in the outcome or associated risks have been associated with performing this intracranial research (*46*).

The behavioral task we employed is a probabilistic reward and punishment learning task with reversal learning where participants’ actions were reinforced or punished with monetary gains or losses. Participants are instructed and actually paid a bonus according to the dollar amounts they earn in the task. Unbeknownst to the participants, the task is setup in stages (fig. S2), such that the initial stage (phase 1) is biased towards probabilistic gain trials (binary outcomes: $1 or $0) where participants can earn an initial reserve of cash before entering phase 2, which introduces trials with probabilistic losses (binary outcomes: -$1 or $0). In the final stage (phase 3), the probabilities of gain or loss outcomes associated with the choice cues are held constant, but the magnitudes of the outcomes are changed, such that the expected values change which options should be expected to pay the most or least (fig. S2; *43*). Optimal performance on this task requires participants to learn from positive and negative feedback to select the option on each trial that maximizes the expected reward and minimizes the expected punishment.

### Human dopamine levels and temporal difference reward prediction errors

Behavioral data demonstrated that participants learned the PRP task’s incentive structure: they chose the best option on a given trial more often than chance (fig. S3). To test whether sub-second dopamine fluctuations in human caudate reflected TD-RPEs, we extracted time series of dopamine levels on each trial aligned to the moments of option presentation, action selection, and outcome presentation, each of which were expected to elicit TD-RPEs during the course of the task. We fit a temporal difference reinforcement learning (TDRL) model to participant behavior and compared the average dopamine timeseries estimates for positive TD-RPEs (n=640) and negative TD-RPEs (n=524) (Fig. 2A,B; fig. S4). We found that, across all trials, sub-second dopamine fluctuations in human caudate did not significantly distinguish positive versus negative TD-RPEs (two-way ANOVA: F_RPE-sign_(1,6) = 1.40, p = 0.24; Fig. 2A). However, separating dopamine responses into reward-versus punishment-trial types revealed that dopamine release distinguished TD-RPEs on reward trials (two-way ANOVA: F_RPE-sign_(1,6) = 5.83, p = 0.016; Fig. 2B) but did not distinguish TD-RPEs on punishment trials (two-way ANOVA: F_RPE-sign_(1,6) = 0.12, p = 0.72; Fig. 2B). Notably, on reward trials, dopamine fluctuations discriminated TD-RPEs within 300ms following a prediction error (one-tailed independent samples t-tests [(RPE>0) > (RPE<0)]: t_200ms_(688) = 2.57, p = 0.0052; t_300ms_(688) = 2.04, p = 0.021).

**Figure 2.**
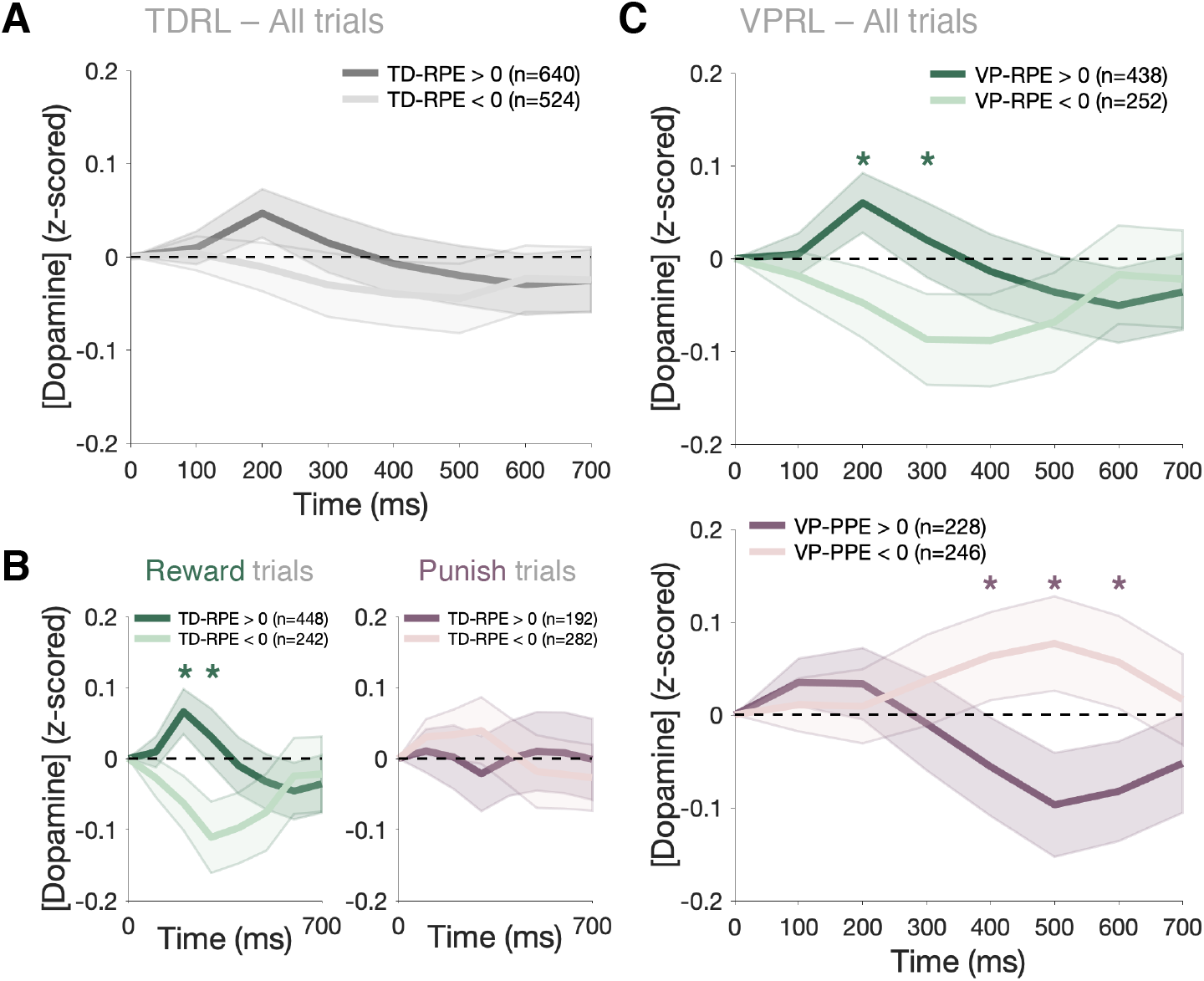
Phasic dopamine levels in human caudate reflect reward and punishment prediction errors. Dopamine responses from 0-700ms following prediction errors across all trials in the PRP task are categorized by prediction error sign and trial type. (**A**) Phasic dopamine transients fail to separate positive and negative TD-RPEs. (**B**) Dopaminergic TD-RPE responses sorted by trial type: gain trials (left panel), loss trials (right panel). (**C**) Phasic dopamine transients across all trials sorted by VP-RPEs sign (top panel) and VP-PPEs sign (bottom panel). Asterisks denote p < 0.05.

### Human dopamine levels and valence-partitioned prediction errors

Prior work demonstrated that dopaminergic responses could track punishment prediction errors (*12, 30–33*), but results shown in Figure 2B suggest that dopamine fluctuations do not reflect *temporal difference reward learning* when the outcome stimulus is punishing (e.g., monetary losses). Thus, we hypothesized that dopamine may encode punishment prediction errors, but as an independent, punishment-specific valuation system (*44*). We tested this hypothesis by fitting to participant behavior a valence-partitioned reinforcement learning (VPRL) model that expresses the independence of reward and punishment learning explicitly (*42, 44*).

Fitting subjects’ behavior to a VPRL model resulted in a better fit to participant behavior compared to TDRL (table S1; *43*). We replicated these results in an independent cohort of healthy human adults (N=42) who completed the PRP task on a computer in a behavioral laboratory setting (*43*; table S1, fig. S3, S5, S6). Further comparisons revealed that VPRL algorithms may perform reward and punishment learning more efficiently than traditional TDRL models that do not partition appetitive and aversive stimuli (fig. S5, S6).

Taken together, our behavioral analyses are consistent with participants adaptively learning the PRP task structure by updating representations of rewarding experiences independently from representations of punishing experiences. Thus, we next tested the hypothesis that dopamine release encoded valence-partitioned RPEs (VP-RPEs) and valence-partitioned punishment prediction errors (VP-PPEs) by sorting dopamine release time series data by the VPRL model-specified prediction errors: positive VP-RPEs (n=438), negative VP-RPEs (n=252), positive VP-PPEs (n=228), or negative VP-PPEs (n=246) (Fig. 2C; fig. S4). We found that dopamine transients distinguished VP-RPEs on reward trials within the same time window as found for TD-RPEs (two-way ANOVA: F_RPE-sign_(1,6) = 3.48, p = 0.06; one-tailed independent samples t-tests [(RPE>0) > (RPE<0)]: t_200ms_(688) = 2.1, p = 0.018; t_300ms_(688) = 1.66, p = 0.049; Fig. 2C). However, we also observed that phasic dopamine responses effectively distinguished VP-PPE signals (two-way ANOVA: F_RPE-sign_(1,6) = 8.08, p = 0.0045; Fig. 2C) within a temporal window distinct from VP-RPE responses, lasting from 400-600ms following a prediction error (one-tailed independent samples t-tests [(PPE>0) < (PPE<0)]: t_400ms_(472) = -1.68, p = 0.047; t_500ms_(472) = -2.3, p = 0.011; t_600ms_(472) = -1.90, p = 0.029). These results demonstrate that sub-second dopamine fluctuations in human caudate may encode valence-partitioned reward and punishment prediction errors.

### Decoding reward and punishment prediction errors from dopamine levels

Fluctuations in extracellular dopamine levels are expected to provide an interpretable signal to downstream neural structures. To determine whether the signals we report (Fig. 2C) are robust enough to be decoded, we trained logistic classifiers to distinguish dopamine time series resulting from positive and negative prediction errors on reward trials (Fig. 3A,B) or positive and negative prediction errors on punishment trials (Fig. 3C,D; *43*). The classifiers trained to discriminate positive versus negative reward prediction errors (TD-RPEs or VP-RPEs on rewarded trials) performed comparably for both TDRL and VPRL models (Fig. 3A,B). Conversely, classifiers trained to discriminate positive from negative punishment prediction errors (TD-RPEs or VP-PPEs on punishment trials) only succeeded when the dopamine time series were parsed according to the VPRL model, and performed at chance level when the dopamine transients were hypothesized to be encoded by TDRL (Fig. 3C,3D).

**Figure 3.**
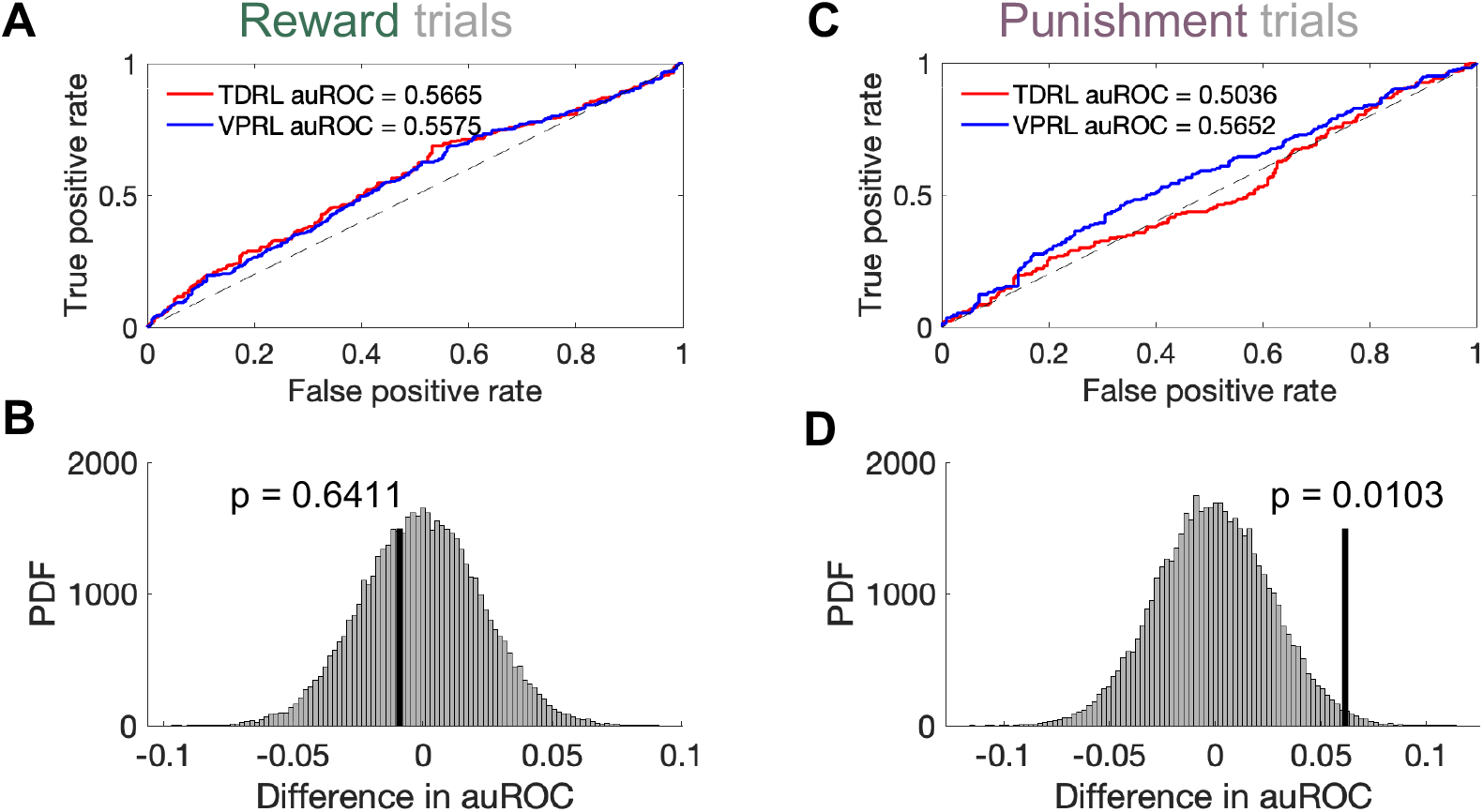
VPRL reward- and punishment-prediction errors can be decoded from human dopamine transients. Performance of the logistic classifier trained on **(A)** TDRL-(red) or VPRL-derived (blue) positive and negative RPEs is comparable across models, with **(B)** the difference in the area under the receiver operating characteristic curve (auROC) values not being statistically significant; p-value derived from permutation test with 50,000 iterations. **(C**,**D)** Same as **(A**,**B)** but for punishment trials; the difference in auROC values for the PPE logistic classifiers were significantly different for TDRL and VPRL (p= 0.0103).

In summary, we demonstrate in humans that sub-second dopamine fluctuations in the caudate nucleus reflect reward and punishment prediction error signals as predicted by a valence-partitioned reinforcement learning framework. Collectively, our results suggest that human decision-making is influenced by independent, parallel processing of appetitive and aversive experiences that can affect modulation of dopamine release in striatal regions on rapid timescales (hundreds of milliseconds). Our findings provide a new perspective on previous reports that dopamine fluctuations in human striatum appear to superpose actual and counterfactual information during risky decision-making (*38, 39*). The results of the present study are consistent with the idea that behavioral reinforcers are processed by independent neural systems according to the valence of the stimulus. Related ideas have been proposed, for instance that rewards and punishments are integrated together during learning (as opposed to being processed independently), leading to a “zero-sum” prediction error that is signaled by dopamine neurons only if the prediction error is positive (i.e., rewarding; *47*); or, that positive and negative RPEs are learned about “asymmetrically” (i.e., different learning rates; *48, 49*). Importantly, however, these proposals are computationally and algorithmically distinct from what is proposed by a model like VPRL where learning about appetitive and aversive stimuli is performed by independent systems prior to being compared to valence-specific expectations (*44*).

There are also a multiplicity of plausible neural mechanisms that could give rise to the present data. Notably, the timing of phasic dopamine response to punishing events is consistent with proposed neuroanatomical circuitry by which aversive stimuli may modulate dopamine neuron activity (*50*). Combining recordings of somatic spiking activity and neurotransmitter release at target brain regions could test more comprehensively, for instance, whether distinct sub-populations of dopamine neurons may be activated to signal valence-specific prediction errors, or whether a separate neural system controls the timing and direction of dopaminergic activity in response to valent behavioral reinforcers.

The approach used to collect the data presented here are severely constrained by the requirement of standard-of-care neurosurgical procedures that provide ‘safe passage’ deep into the human brain. Significant challenges lay ahead for future efforts to determine whether these findings generalize to other brain regions and other patient populations, including neurologically healthy humans; however, these data demonstrate that such recordings are feasible. A growing number of conditions use DBS to effect symptom management, including Parkinson’s disease, substance use disorders, and depression. Together, patient volunteers from these and other populations working with clinical research teams can provide significant insight into human brain function, human experience, and the mechanisms in human neural systems that are altered in human brain disorders.

## Acknowledgments

We thank the participants of this study and the research and surgical nursing staff at Atrium Health Wake Forest School of Medicine for their support and cooperation.

## Funding

This work was supported by National Institutes of Health grants R01MH121099 (KTK), R01DA048096 (KTK), R01MH124115 (KTK), P50DA006634 (KTK), 5KL2TR001420 (KTK), F31DA053174 (LPS), T32DA041349 (LPS), and F30DA053176 (BL).

## Author contributions

L.P.S. designed and performed behavioral and FSCV data analysis, interpreted results, wrote and edited original manuscript drafts, approved final manuscript; A.J. coded behavioral tasks, collected data, analyzed FSCV data, edited and approved final manuscript; B.L. collected data, edited and approved final manuscript; E.D. collected data, edited and approved final manuscript; A.W.L. conceived of surgical strategies for safe and effective placement of electrodes for human FSCV, performed surgical placement of electrodes for human FSCV experiment, collected data, edited and approved final manuscript; S.B.T. conceived of surgical strategies for safe and effective placement of electrodes for human FSCV, performed surgical placement of electrodes for human FSCV experiment, collected data, edited and approved final manuscript; P.R.M. interpreted results, edited and approved final manuscript; K.T.K. conceived the study, designed experiments, supervised and guided data collection and analysis, interpreted results, wrote and edited original manuscript drafts, and approved final manuscript.

## Competing interests

The authors declare no competing interests.

## Data and materials availability

Anonymized individual-level participant behavioral task data and neurochemical time series data used in this study may be made available upon submission of a formal project outline from any qualified investigator to the corresponding author and subsequent approval by the corresponding author in line with data protection regulations of Wake Forest University School of Medicine Institutional Review Board (IRB). Custom-written analysis scripts for generating the behavioral and neurochemical time series results of this manuscript are maintained in a private github repository (*insert link upon acceptance*) that may be shared upon request from any qualified investigator to the corresponding author.

## Materials and Methods

### Patient recruitment and informed consent

A total of 11 patients (6 female, 5 male, age range = 48-82, mean age +/-SD = 67.5 +/-10.9) diagnosed with essential tremor (ET) and approved candidates for DBS treatment participated in this study. Three patients performed the procedure while carbon fiber microelectrodes recorded dopamine release in their caudate, and one patient performed the procedure while a carbon fiber microelectrode recorded dopamine release in their thalamic ventralis intermediate nucleus (VIM). The other seven patients performed the task while recordings were made with a tungsten microelectrode. All eleven patients’ behavioral data were included in analyses for hierarchical parameter estimation; however, the tungsten microelectrode (n=7) and thalamic VIM (n=1) neurochemical recordings are not presented in the present work. After informed written consent was obtained from each patient, they were given details about the decision-making task (i.e., probabilistic reward and punishment task) and were familiarized with the type of outcomes experienced during game play and the controllers used for submitting responses. The experiment was approved by the Institutional Review Board (IRB#: IRB00017138) of Wake Forest University Health Sciences (WFUHS). Out of the eleven patients that participated in the study, four patients did not complete all 150 trials of the task (range = 121-148 trials).

In addition to the cohort of ET patients, a behavior-only cohort of healthy adult humans (N=42; 19 female) was recruited from the local Winston-Salem community to complete the PRP task. Informed written consent was obtained from each participant, and the experiment was approved by the Institutional Review Board (IRB#: IRB00042265) of Wake Forest University Health Sciences (WFUHS). All behavioral experiments were conducted at WFUHS.

### Probabilistic Reward and Punishment (PRP) task experimental procedure

The PRP task (**Fig. 1A; fig. S2**) is a 150-trial, two-choice monetary reward and punishment learning task, where chosen options are reinforced probabilistically with either monetary gains (or no gain) or monetary losses (or no loss). Six options (represented by fractal images) comprise the set of possible actions, with each option assigned to one of three outcome probabilities (25%, 50%, and 75%) and one of two outcome valences (monetary gain or loss); thus, there are three reward-associated ‘gain/no gain’ options and three ‘loss/no loss’ options in the task, and the assignment of options to outcome probabilities and valences is randomized across participants. On each trial, two out of the six options are presented (note that option pairings are random, not fixed); depending on the phase of the task (Phase 1: trials 1-25; Phase 2: trials 26-75; Phase 3: trials 76-150), either two of the three ‘gain/no gain’ options are presented (i.e., ‘gain/no gain’ trials), or two of the three ‘loss/no loss’ options are presented (i.e., ‘loss/no loss’ trials), or one of each ‘gain/no gain’ and ‘loss/no loss’ options are presented (i.e., ‘mixed’ trials). Participants were told that certain options in the PRP task would earn them money and some options would lose them money, and participants were instructed that their goal was to maximize their earnings on the task and that they would receive their total earnings as a bonus monetary payment at the end of the study visit.

At the beginning of the experiment (Phase 1, trials 1-25), each trial starts with the presentation of two of the three possible ‘gain/no gain’ options, and participants are reinforced with either a monetary gain or nothing ($1 or $0) according to the chosen option’s fixed probability. In Phase 2 (trials 26-75), the task introduces ‘loss/no loss’ trials which present two of the three ‘loss/no loss’ options that result in either a monetary loss or nothing (-$1 or $0) with fixed probabilities. In this phase, there are an equal number of ‘gain/no gain’ and ‘loss/no loss’ trials, randomly ordered. In Phase 3 (trials 76-150), two options are presented randomly such that any trial may consist of two ‘gain/no gain options, two ‘loss/no loss’ options, or one ‘gain /no gain and one ‘loss/no loss’ option. Moreover, in Phase 3 the outcome magnitudes of all options change such that the 25%, 50%, and 75% ‘gain’ options now payout $2.50, $1.50, and $0.50, respectively, and the 25%, 50%, and 75% ‘loss’ options now lose -$1.25, -$0.75, and -$0.25, respectively (see dashed lines in **fig. S2**).

On each trial, participants select an option at their own pace. Once a selection has been made, the unchosen option disappears at the same time that the chosen option is highlighted, and this screen lasts for three seconds. The outcome is then displayed for one second followed by a blank screen that lasts for a random time interval (defined by a Poison distribution with λ = 3 seconds) before the next trial begins. Additionally, on each trial with probability 0.33, the blank screen following the outcome presentation is followed by a subjective feeling rating screen that consists of the text “How do you feel about the last outcome?” and a visual-analog rating scale with a vertical bar cursor that can be moved by the participant. Participants are asked to rate their feelings about the experienced outcome with this visual-digital scale, after which the blank screen reappears for another random time interval before a new trial begins.

### Behavioral data analysis

#### Temporal Difference Reinforcement Learning model

In the standard TDRL model (*14,51*), the expected value of a state-action pair Q(*s*_*i*_, *a*_*i*_), where *i* indexes discrete time points in a trial, is updated following selection of action *a*_*i*_ in state *s*_*i*_ according to:

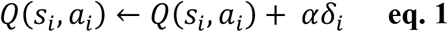

where 0 < α < 1 is a learning rate parameter that determines the weight prediction errors have on updating expected values, and *δ*_*i*_ is the TD reward prediction error term:

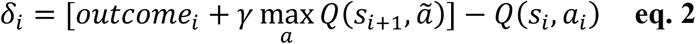

where *outcome*_*i*_ is the outcome (positive or negative) experienced in state *s*_*i*_ after taking action *a*_*i*_, 0 < *γ* < 1 is a temporal discount parameter that discounts outcomes expected in the future relative to immediate outcomes, and max Q(*s*_*i*+1_, *a*>) is the maximum expected action value over all actions *a*> afforded in the next state *s*_*i*+1_. We defined the trials of the PRP task as consisting of *i* = {1, 2, 3, 4} event time points (1: options presented; 2: action taken; 3: outcome presented; 4: (terminal) transition screen). We modeled participant choices (*choice*_*t*_) on each trial *t* of the PRP task with a softmax choice policy (i.e., categorical logit choice model) that assigns probability to choosing each of the two options presented on a trial according to the learned Q-values of the two options. For example, for a trial that presents option 2 and option 5, the corresponding action values at the moment of option presentation, Q(*s*_1_, op*t*_2) and Q(*s*_1_, op*t*_5), are used to compute the probability of selecting each option:

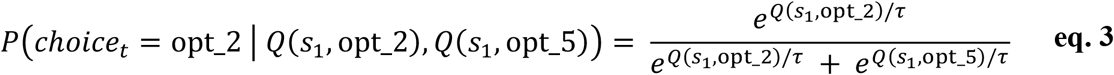

where 0 < τ < 20 is a choice temperature parameter that determines the softmax function slope and parameterizes an exploration versus exploitation trade-off where higher temperature values lead to a more randomized choice selection policy and lower temperature values lead to a more winner-take-all, deterministic choice policy.

#### Valence-Partitioned Reinforcement Learning (VPRL) model

For valence-partitioned RL (VPRL, *44*), we extend the standard TDRL framework by specifying that two separate value representations are learned for each action, corresponding to the rewarding value and punishing value of each action, and that separate (neural) systems signal reward- and punishment-specific prediction errors to update the reward- and punishment-associated action values, respectively. In this way, VPRL treats ‘Positive’ (*P*) and ‘Negative’ (*N*) outcomes as though separate, parallel *P*- and *N*-systems effectively establish a partition between the processing of rewarding and punishing outcomes. *P*- and *N*-system action values are estimated (*Q*^*p*^ and *Q*^*N*^, respectively) independently, though each system learns these outcome valence-specific action values using temporal difference learning (see **eqs. 4-7**). We model the integration of *Q*^*p*^ and *Q*^*N*^ in the simplest manner (i.e., subtraction; **eq. 8**) when value-based decisions must be made, though alternative approaches for integrating these value estimates may be investigated in future work.

In VPRL, *P-* and *N-*systems update action value representations via TD-prediction errors on every episode, but by valence-specific rules (*P-system:* **eq. 4**; *N-system:* **eq. 5**). The *P-*system only tracks rewarding (i.e., appetitive) outcomes (*outcome*_*i*_ > 0, **eq. 4**) and the *N-*system only tracks punishing (i.e., aversive) outcomes (*outcome*_*i*_ < 0, **eq. 5**); both systems encode the opposite-valence outcomes and null outcomes as though no outcome occurred.

Thus, For the *P*-system, the reward-oriented TD prediction error is:

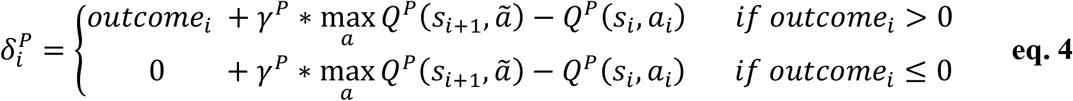

where 0 < *γ*^p^ < 1 is the *P-*system temporal discounting parameter (directly analogous to the standard TDRL temporal discounting parameter).

The *N-*system similarly encodes a punishment-oriented TD prediction error term:

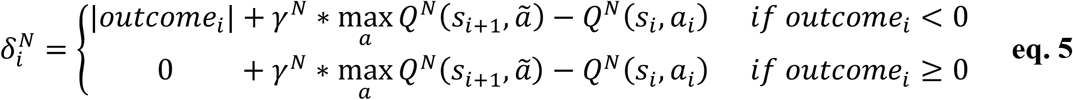

where 0 < *γ*^4^ < 1 is the *N-*system temporal discounting parameter and |*outcome*_*i*_| indicates the absolute value of the (punishing) outcome. We use the absolute value of the outcome so that the *N*-system positively communicates punishments of varying magnitudes, reflecting a neural system that increases its firing rate for larger-than-expected punishments and decreases its firing rate for smaller-than-expected punishments.

The *P-* and *N-*systems prediction errors update expectations of future rewards or punishments of an action, respectively, according to the standard TD learning update rule but for each system independently:

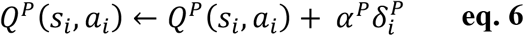

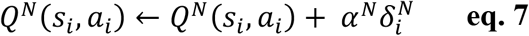

where 0 < *α*^*p*^ < 1 and 0 < α^N^ < 1 are learning rates for the *P-* and *N-*systems, *Q*^*p*^(*s*_*i*_, *a*_*i*_) is the expected state-action value learned by the *P-*system, and *Q*^*N*^(*s*_*i*_, *a*_*i*_) is the expected state-action value learned by the *N-*system.

We compute a composite state-action value for each action by contrasting the *P-* and *N-* system Q-values,

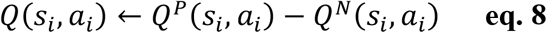

which is entered into the categorical logistic choice model (e.g., softmax policy, **eq. 3**) as for the TDRL model above.

#### Alternative reinforcement learning models

Apart from the TDRL and VPRL models described above, we fit ‘asymmetric’ versions of these models to participant choice behavior on the PRP task. ‘Asymmetric’ TDRL and VPRL models are defined by using distinct learning rate parameters for prediction errors that are positive or negative. For asymmetric TDRL, this amounts to changing **eq. 1** to:

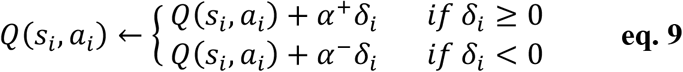

where 0 < α^+^ < 1 is the learning rate for positive TD-RPEs and 0 < α^5^ < 1 is the learning rate for negative TD-RPEs; the rest of the traditional TDRL model remains the same. For asymmetric VPRL, **eq. 6** and **eq. 7** are changed to:

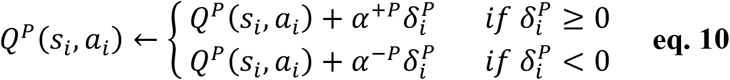

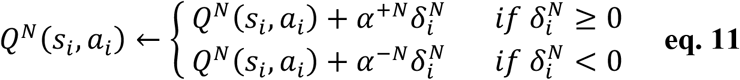

where 0 < α^+p^, α-^p^ < 1 are learning rate parameters for positive and negative VP-RPEs, respectively, and 0 < α^+N^, α-^N^ < 1 are learning rate parameters for positive and negative VP-PPEs, respectively; the rest of the original VPRL model remains the same.

#### Reinforcement learning hierarchical model parameterization

We specified a hierarchical structure to all computational models to fit participant choice behavior on the PRP task. Individual-level parameter values are drawn from group-level distributions over each model parameter. This hierarchical modeling approach accounts for dependencies between model parameters and biases individual-level parameter estimates towards the group-level mean, thereby increasing reliability in parameter estimates, improving model identifiability, and avoiding overfitting (*52*). These hierarchical models therefore cast individual participant parameter values as deviations from a group mean.

Formally, the joint posterior distribution *P*(*ϕ*, θ|*y, M*_*i*_) over group-level parameters ϕ and individual-level parameters θ for the *i*-th model *M*_*i*_ conditioned on the data from the cohort of participants *y* takes the form

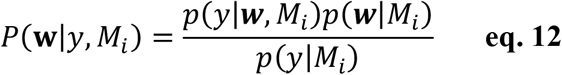

where we simplify the notation to *P*(***w***|*y, M*_*i*_), with ***w*** = {ϕ, θ}) being a parameter vector consisting of all group- and individual-level model parameters for model *M*_*i*_. Here, *P*(*y*|***w***, *M*_*i*_) is the likelihood of choice data *y* conditioned on the model parameters and hyperparameters, *P*(*y*|*M*_*i*_) is the marginal likelihood (model evidence) of the data given a model, and *P*(***w***|*M*_*i*_) is the joint prior distribution over model parameters as defined by the model *M*_*i*_, which can be further factorized into the product of the prior on individual-level model parameters conditioned on the model hyper-parameters, *P*(θ|ϕ, *M*_*i*_), times the prior over hyper-parameters *P*(ϕ|*M*_*i*_). We define the prior distributions for individual-level model parameters (e.g., θ_*TDRL*_ = {α, τ, γ} for *M*_*i*_ = TDRL) and the hyper-priors of the means −∞ < μ_(.)_ < +∞ and standard deviations 0 < σ_(.)_ < +∞ of the population-level parameter distributions (e.g., ϕ_*TDRL*_ = {μ _α_ μ_τ_, μ_γ_ σ_α_ σ_τ_, σ_γ_}) to be standard normal distributions. We estimated all parameters in unconstrained space (i.e., −∞ < μ_γ_ < +∞) and used the inverse Probit transform to map bounded parameters from unconstrained space to the unit interval [0,1] before scaling parameter estimates by the parameter’s upper bound:

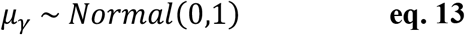

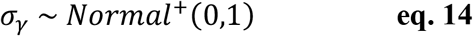

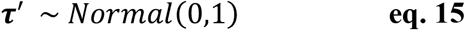

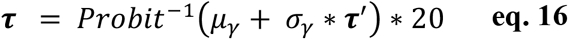

where bold terms indicate a vector of parameter values over participants. This non-centered parameterization (*53*) and inverse Probit transformation creates a uniform prior distribution over individual-level model parameters between specified lower and upper bounds. Note that for learning rate and temporal discount parameters, the scaling factor (upper bound) was set to 1, whereas it was set to 20 for the choice temperature parameter. We used the Hamiltonian Monte Carlo (HMC) sampling algorithm in the probabilistic programming language Stan (*54*) via the R package *rstan* (v. 2.21.2) to sample the joint posterior distribution over group- and individual-level model parameters for both cohorts individually and for all participants combined into a single cohort. For all models and each cohort, we executed 12,000 total iterations (2,000 warm-up) on each of 3 chains for a total of 30,000 posterior samples per model parameter. We inspected chains for convergence by verifying sufficient chain mixing according to the Gelman-Rubin statistic 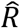, which was less than 1.1 for all parameters.

#### Reinforcement learning model comparison

We compared the fit of each model to participant choice behavior on the PRP task according to their model evidence (i.e., Bayesian marginal likelihood), which represents the probability or ‘plausibility’ of observing the actual PRP task data under each model (*55*). In Bayesian model comparison, the model with the greatest posterior model probability *p*(*M*_*i*_|*y*) is deemed the best explanation for the data *y* and is computed by:

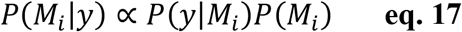

where *P*(*y*|*M*_*i*_) is the model marginal likelihood (i.e., ‘model evidence’), the normalizing constant of **eq. 12**, and *P*(*M*_*i*_) is the model’s prior probability. The model evidence is defined as:

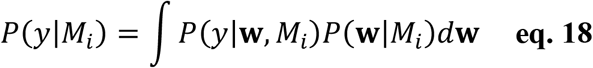

where *P*(***w***|*M*_*i*_) is the prior probability of a model *M*_*i*_’s parameters ***w*** before observing any data and *P*(*y*|***w***, *M*_*i*_) is the likelihood of data *y* given a model and its parameters.

Importantly, the marginal likelihood for each model as defined in **eq. 18** is an optimal measure for performing model comparison as it represents the balance between the fit of each model to the cohort’s data (as captured by the first term in the integral) and the complexity of each model (as captured in the second term of the integral), integrated over all sampled model parameter values. In effect, although more complex or flexible models (i.e., more parameters) are able to predict a greater variety of behaviors and therefore increase the data likelihood *P*(*y*|***w***, *M*_*i*_), more complex models have a higher dimensional parameter space and therefore must necessarily assign lower prior probability to the parameter values *P*(***w***|*M*_*i*_). In this way, the marginal likelihood of a model automatically penalizes model complexity, sometimes referred to as the ‘Bayesian Occam razor’ (*55*).

To compare the TDRL and VPRL models (i.e., *M*_*i*_ and *M*_2,_respectively), the relative posterior model probability can be defined as:

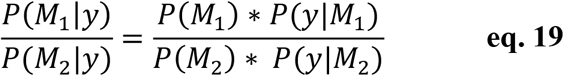

where the ratio of posterior model probabilities 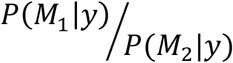 is referred to as the “posterior odds” of TDRL relative to VPRL; *P*(*M*_*i*_) and *P*(*M*_2_) are the prior probabilities of the TDRL and VPRL models, respectively; and the ratio of marginal likelihoods 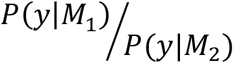 is termed the “Bayes factor”, which is a standard measure for Bayesian model comparison. By assigning equal prior probabilities over the set of candidate models, each model’s evidence *P*(*y*|*M*_*i*_) can be used to rank each model in the set for comparison. The marginal likelihoods are computed as log-scaled and therefore the Bayes factor is computed as the difference between log marginal likelihoods for two models; a positive value for the Bayes factor indicates greater support for *M*_*i*_ (the model in the numerator of **eq. 19**), whereas a negative value for the Bayes factor indicates greater support for *M*_2_. We estimated the log model evidence (marginal likelihood) for all models for each cohort, and for all participants combined into a single cohort, using an adaptive importance sampling routing called bridge sampling as implemented in the R package *bridgesampling* (v. 1.1-2; *56*). Bridge sampling is an efficient and accurate approach to calculating normalizing constants like the marginal likelihood of models even with hierarchical structure and for reinforcement learning models in particular (*56*). To further ensure stability in the bridge sampler’s estimates of model evidence, we performed 10 repetitions of the sampler and report the median and interquartile range of the estimates of model evidence. The model with the maximum (i.e., least negative) model evidence is the preferred model.

In addition to the standard Bayesian model comparison using model marginal likelihoods, we estimated each model’s Bayesian leave-one-out (LOO) cross-validation predictive accuracy, defined as a model’s expected log predictive density (ELPD-LOO; *57*):

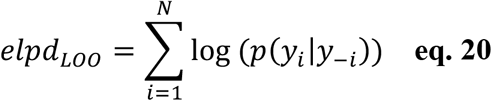

where the posterior predictive distribution *p*(*y*_*i*_|*y*_5*i*_) for held-out data *y*_*i*_ given a set of training data *y*_5*i*_, is

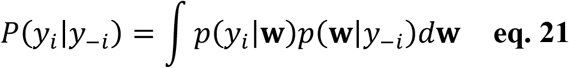

The ELPD is an estimate of (i.e., approximation to) the cross-validated accuracy of a given model in predicting new (i.e., held-out) participant data, given the posterior distribution over model parameters fit to a training set of participant data (*57*). We approximate this integral via importance sampling of the joint posterior parameter distribution given the training data *p*(***w***|*y*_5*i*_) using the R package *loo* (v. 2.3.1; *57*).

We repeated this model comparison analysis (**table S1**) for the behavior-only cohort and a ‘meta-analytic’ cohort combining the ET patients and behavioral participants (N=53). Running the model comparison analysis in triplicate allowed us to assess the replicability of the model comparison results, and employing multiple model comparison criteria allowed us to assess the robustness and generalizability of the model comparison results. We elected to focus the subsequent behavioral and neurochemical analyses on the basic TDRL and VPRL models since the computational differences between these models most directly address the neurobiological mechanism that was our main target of investigation: the partitioned signaling of reward and punishment prediction errors; all subsequent behavioral analyses and neurochemical time series analyses of the ET cohort used the computational model fits to the ET cohort alone.

#### Model and parameter recovery

We performed a model recovery analysis to validate that our Bayesian model comparison analysis is able to accurately identify the true generative model of choice behavior on the PRP task. For this model recovery analysis, we simulated choice behavior on the PRP task for both the ET (N=11) and behavioral (N=42) cohorts using the mean individual-level parameter values for TDRL and VPRL models and then computed model comparison criteria for the TDRL and VPRL models to determine whether the model comparison analysis identified the true generative model as the best model (**table S2**).

To validate that our hierarchical computational model fitting procedure is able to accurately estimate model parameters for each participant and for TDRL and VPRL models, we performed a parameter recovery analysis. We determined whether the empirical parameter distributions for both cohorts were credibly different by computing the difference between the ET and behavioral cohorts’ group-level TDRL and VPRL parameter distributions, which revealed no credible differences in any TDRL or VPRL model parameter between the cohorts (**fig. S7**). Given this result, and since the larger sample size in the behavioral cohort increases the robustness of the parameter recovery analysis results, we elected to perform the parameter recovery analysis using the behavioral cohort’s data. We first calculated the mean TDRL and VPRL parameter values for each participant in the behavioral cohort to simulate choice data sets (N=42) on the PRP task (using new option presentation sequences), re-fitted the TDRL and VPRL models to the simulated PRP data set, and then computed the Pearson’s correlation coefficient between the mean model parameters fitted to the actual participant PRP data and the simulated PRP data.

#### Electrochemistry data analysis

##### General description of human voltammetry approach

The human fast-scan cyclic voltammetry (FSCV) protocol used in the current study has been extensively described in previous publications (*38–41*), and therefore we give a brief general description here. The human voltammetry protocol, which involves the construction of custom carbon-fiber microelectrodes for use in the human brain (*38,40*), is designed as a human-level extension of traditional voltammetry protocols used in model organism (e.g., rodent) and *ex-vivo* slice or culture preparations. The specific electrochemical properties of the custom electrodes used in the human voltammetry protocol have been validated in the rodent brain as matching those of rodent electrodes (*40*). Additionally, the voltage waveform and cycling frequency of the stimulating current, as well as the sampling rate of the current time series during the voltage sweeps used in the human protocol, are identical to those used in rodent studies (*26*).

The central difference between the human voltammetry protocol used here (*38, 39, 41*) and traditional voltammetry protocols is the statistical method employed to estimate the *in-vivo* concentration of different neurochemical analytes. Specifically, in traditional voltammetry protocols, estimating the concentration of an analyte of interest (e.g., dopamine) involves performing principal components regression on recorded currents (voltammograms), wherein the principal component time series used as regressors are derived from an *in-vitro* data set of voltammograms of known concentrations of the analyte of interest. By contradistinction, the statistical method used for analyte concentration estimation in the human voltammetry protocol adopts a supervised statistical learning approach. This approach involves training an elastic net-penalized linear regression model on *in-vitro* voltammograms of known concentrations of analytes of interest (e.g., dopamine, serotonin), varying levels of pH, and common metabolites of target analytes (e.g., DOPAC, 5-HTIAA) or other neurotransmitters (e.g., norepinephrine; *58*). In this protocol, multiple carbon-fiber microelectrodes identical to those used for human recordings were used to collect the *in-vitro* training datasets, and the penalized linear regression model is optimized via cross-validation to reduce the out-of-probe error. This penalized cross-validation procedure has the added benefits of reducing bias in model performance due to overfitting on training data and automatically selecting and regularizing model coefficient values (via the elastic net), thereby providing reliable estimation performance when recovering analyte concentrations from the electrodes used during the human voltammetry experiments. This approach provides more reliable estimates of dopamine than principal components regression (*38*), especially under different pH levels. Additionally, this approach reliably and accurately differentiates mixtures of dopamine and serotonin from a background of varying pH (*39, 41*) and changing levels of dopamine or serotonin metabolites or other neurochemical species like norepinephrine (*58*).

##### FSCV carbon-fiber microelectrodes and experimental protocol

The FSCV protocol as well as the construction of carbon-fiber microelectrode probes and the specifications of the mobile electrochemistry recording station have been extensively described in previous work (*38, 40*). Briefly, custom carbon-fiber microelectrodes for human FSCV experiments were placed in the caudate nucleus as determined by DBS surgery planning for ET patients. We note that electrode placement within the caudate nucleus is different for each patient in accordance with the patient-specific trajectory of the DBS electrode used for treatment. The FSCV protocol consisted of an equilibration phase and an experiment phase where the voltammetry measurement waveform – a triangular waveform starting at -0.6 V, ramping up to a peak of +1.4 V at 400 V/s, and ramping back down to -0.6 V at -400 V/s – was first cycled at 60 Hz for 10 minutes to allow for equilibration of the electrode surface followed by a 10 Hz application of the waveform for the duration of the experimental window encompassing the behavioral task. All recordings of the measurement waveform-induced currents (voltammograms) were collected at a 100 kHz sampling rate.

##### in-vitro training data protocol and neurochemical concentration estimation model training

The *in-vitro* data collected to train the dopamine concentration estimation model consisted of a population of 5 carbon-fiber microelectrodes identical to those used in the human voltammetry experiments. Each probe contributed 16 datasets (one per solution mixture), with each dataset consisting of 2 minutes’ worth of voltammogram recordings in mixture solutions of known concentrations of dopamine, DOPAC, and ascorbic acid (from 0-1500nM in 100nM increments), with a background of varying pH levels (from 7.2-7.6 in 0.1 increments). All voltammograms in the training datasets were sampled at 250 kHz (resulting in 2500 samples per voltammogram) and then downsampled by averaging every 15 samples. The voltammograms used to train the dopamine concentration estimation model were taken over the last 90 seconds of a probe’s 2-minute recording in a given solution, as these later timepoints are less affected by flow or electrode equilibration artifacts that occur in the beginning of recording periods. Each probe therefore contributed a total of 900 voltammograms per each of 16 solution mixtures resulting in a total of 14,400 labeled samples per probe, each corresponding to the probe’s response to mixed levels of dopamine, DOPAC, ascorbic acid, and pH.

Using this *in-vitro* training data set, we fit a penalized linear regression model using the elastic net algorithm (*59*) to predict known concentrations of each analyte, optimized using 10-fold cross-validation. In this model, the target variable (*y*) is an *N*-by-4 matrix of known levels of dopamine, DOPAC, ascorbic acid, and pH, with *N* = 12,960 samples (9/10ths of the 14,400 total samples, with 1/10 held-out for cross-validation); the predictor variable matrix (*x*) is an *N*-by-498 matrix of the corresponding raw and differentiated voltammograms (167 time points per down-sampled voltammogram, plus 166 time points for its first derivative and 165 time points for its second derivative). The linear model coefficients (β) are determined by minimizing the residual sum of squares, subject to the elastic net penalty (*59*):

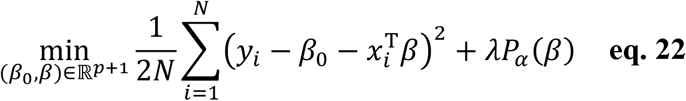

where λ is a penalty term that weighs the influence of the elastic net penalty, *P*_α_ (β):

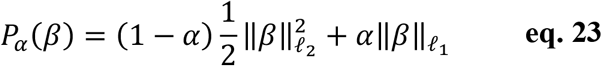

where 0 < α < 1 parameterizes the relative weighting between the ridge (ℓ_2_-norm) and lasso (ℓ_1_-norm) regularizations. The optimal values of β, λ, and α are determined using a 10-fold cross-validation procedure via the *cvglmnet* function of the *glmnet* package in MATLAB. Here, we fixed α = 1 and used the smallest lambda value to estimate dopamine concentrations from *in vivo* experimental recordings.

##### Dopamine time series analysis

Time series of dopamine concentrations for each participant were generated from the optimized elastic net linear regression model with 100 millisecond temporal resolution. We first cut out individual trials’ time series from 1 second (10 samples) before the trial’s option presentation screen to 100 milliseconds (1 sample) before the next trial’s option presentation, z-scored the dopamine concentrations within each trial, and smoothed the within-trial dopamine time series using a 0.3 second (3 sample) sliding-window lagging average (*41*). From these individual trial time series, we extracted individual event-related dopamine responses lasting from 0-700 milliseconds following option presentation, action selection, and outcome presentation. Parametric statistical testing consisted of performing either two-way ANOVA tests (prediction error sign, time) of dopamine responses following all events (**Fig. 2**) or independent samples t-tests at single time points to compare dopamine responses to positive and negative reward and punishment prediction errors following all events (**Fig. 2**). Non-parametric statistical testing (**fig. S4**) consisted of conducting 50,000 permutation tests where we computed the mean difference in dopamine levels in response to positive and negative RPEs and PPEs at each time point and computed p-values as the percentage of permuted mean difference measures that were greater than the absolute value of the actual mean difference.

##### Dopamine prediction error ROC decoding analysis

For the receiver operating characteristic (ROC) analysis (**Fig. 3**), we trained logistic regression models on segments of event-related dopamine fluctuations to classify positive and negative reward and punishment prediction errors. We trained separate classifiers using either TDRL or VPRL computational model-defined fluctuations; that is, the event-related dopamine signals used to train each classifier differed according to whether TDRL and VPRL models specified an event as being either a positive or negative RPE or PPE. For the RPE classifiers, we trained the logistic models for TDRL and VPRL using samples from 200-300ms of the dopamine fluctuations; for the PPE classifiers, we used samples from 400-600ms of the dopamine fluctuations. These RPE- and PPE-specific temporal windows were chosen based on our findings from the dopamine time series analysis (**Fig. 2**). From the fitted classifiers, we computed the area under the ROC curve (auROC) separately for the TDRL- and VPRL-based classifiers using the *perfcurve* function in MATLAB. We compared the relative performance of the TDRL and VPRL classifiers for decoding positive and negative RPEs and PPEs using a permutation test where we computed the difference in auROC values across 50,000 iterations and compared the true auROC values to the permutation test null distribution to obtain p-values.

**Fig. S1.**
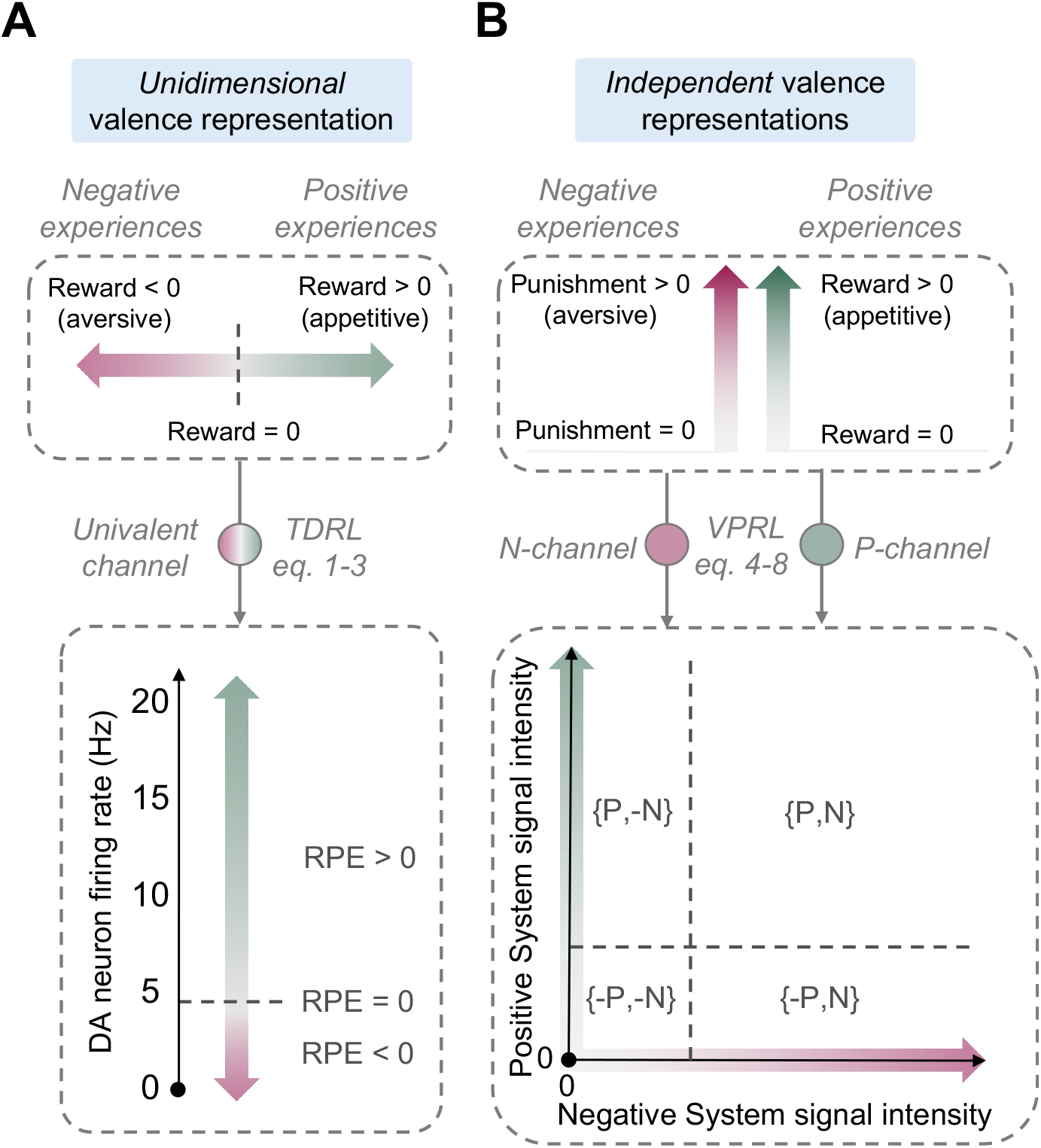
Alternative computational theories of valence processing in reinforcement learning. **(A)** Traditional temporal difference reinforcement learning (TDRL) theory represents rewards and punishments unidimensionally as opposite ends of a single continuous valence dimension. The physiological support for this traditional view is limited by how dopamine neurons might encode aversive outcomes. **(B)** A valence-partitioned reinforcement learning (VPRL) approach instead specifies that rewards and punishments are processed by independent valence-processing systems in parallel. The space spanned by the activity within these two systems of VPRL (the Positive system and Negative system {±P, ±N} space) captures all combinations of possible valent experiences.

**Fig. S2.**
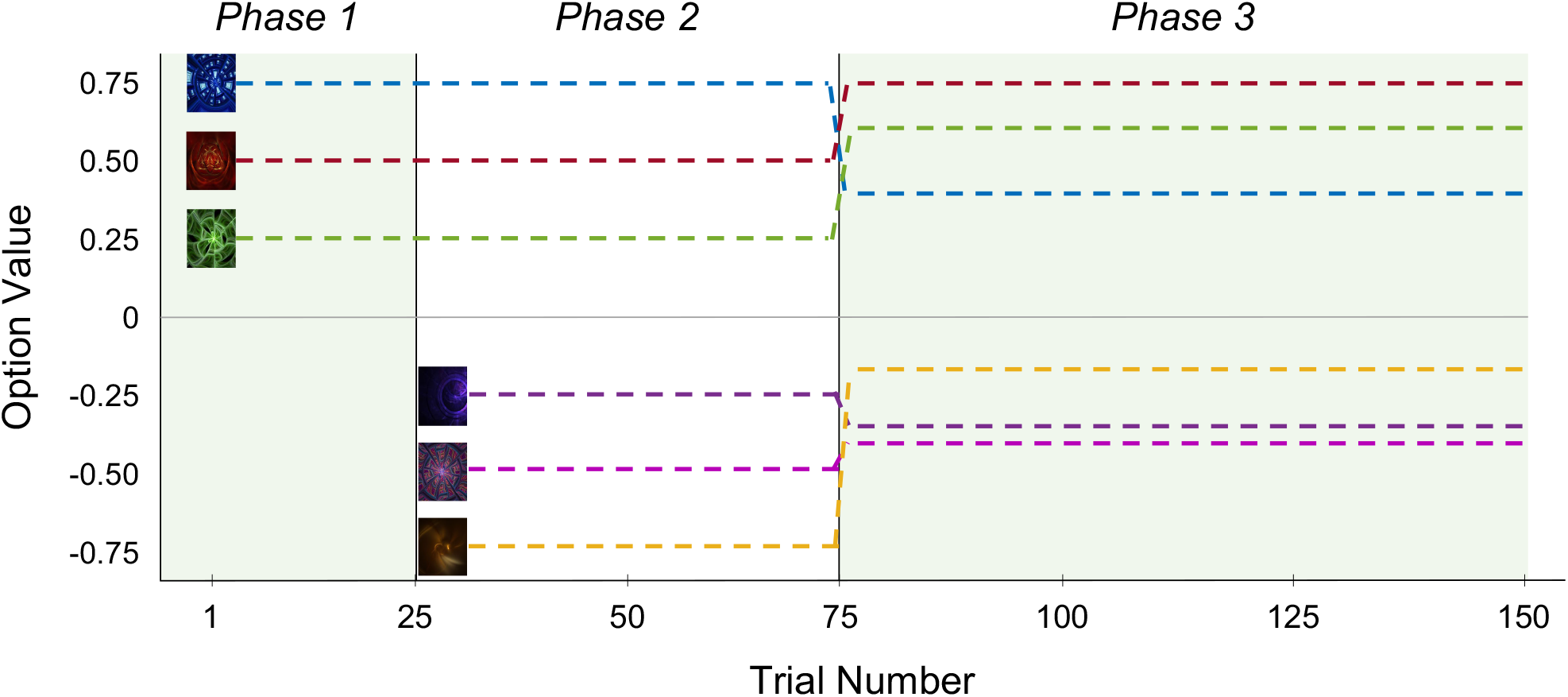
Probabilistic reward and punishment task incentive structure. Depiction of the true expected value of the six options throughout the three phases of the task. Each option has either a low (25%), medium (50%), or high (75%) probability of gaining or losing money; icon-to-probability and icon-to-outcome-valence mappings are randomized across participants. In phase 1, trials 1-25, participants see two of three ‘gain/no-gain’ options, which give binary monetary gains ($0 or $1) according to fixed probabilities; the blue icon corresponds to the 75% gain option (value = $0.75), the brown icon corresponds to the 50% gain option (value = $0.50), and the green icon corresponds to the 25% gain option (value = $0.25). In phase 2, trials 26-75, participants either see two of the three ‘gain/no-gain’ options or two of three ‘loss/no-loss’ options, which give binary monetary losses ($0 or -$1), and there are an equal number of ‘gain/no gain’ and ‘loss/no loss’ trials (i.e., 25 each); the purple icon corresponds to 25% loss (value = -$0.25), the pink icon corresponds to 50% loss (value = -$0.50), and yellow icons correspond to 75% loss (value = -$0.75). In phase 3, trials 76-150, the reversal occurs wherein the outcome magnitude of every option changes (the associated probabilities remain the same); participants see any combination of two of the six options in phase 3. In phase 3, the 75% gain icon now gives binary $0/$0.50 returns (value = $0.38), the 50% gain icon gives $0/$1.50 returns (value = $0.75), and the 25% gain icon gives binary $0/$2.50 returns (value = $0.63); the 75% loss icon now gives binary $0/-$0.25 returns (value = -$0.19), the 50% loss icon gives $0/-$0.75 returns (value = -$0.38), and the 25% loss icon gives $0/-$1.25 returns (value = -$0.31). There are no fixed pairings of options in the task.

**Fig. S3.**
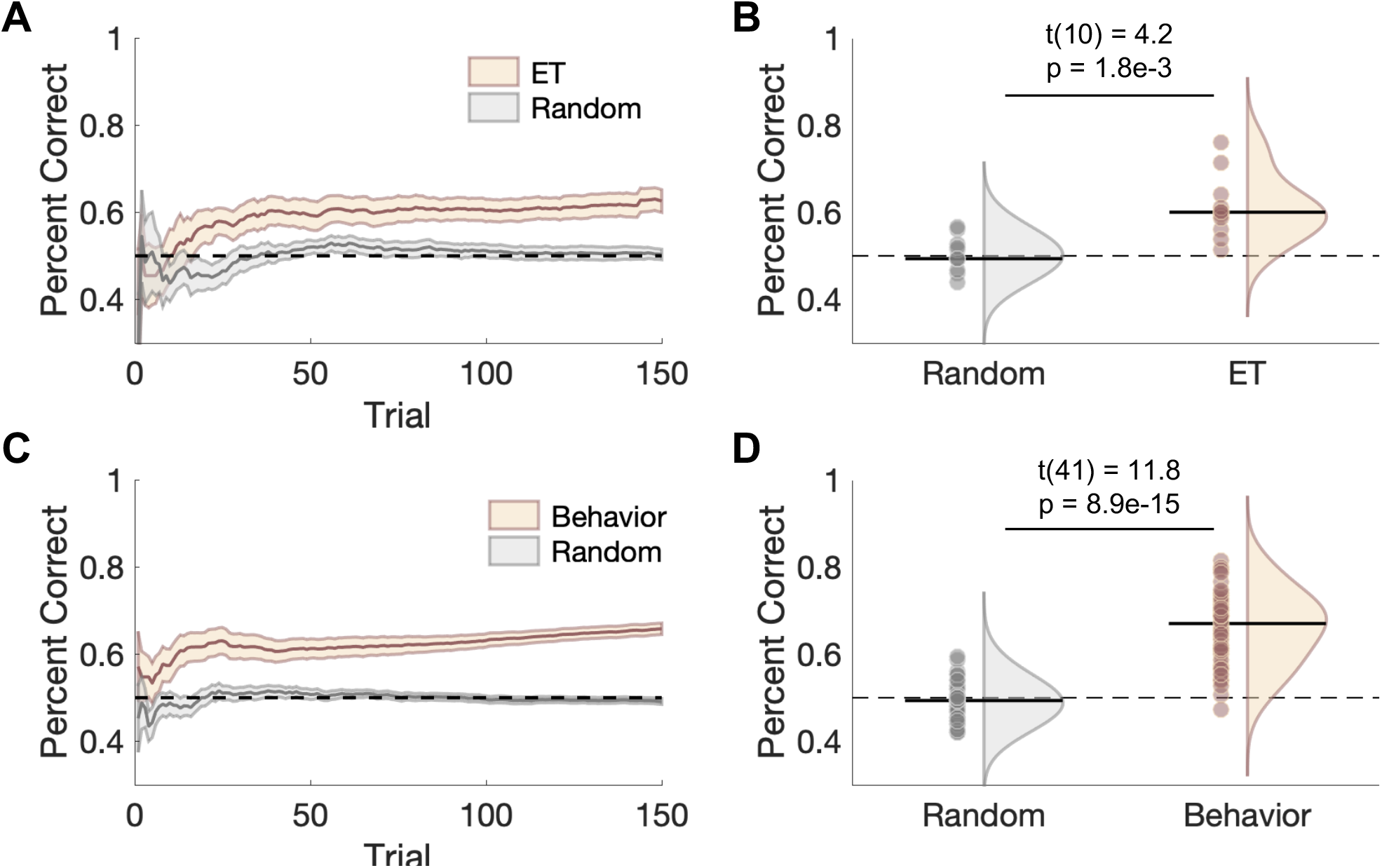
Descriptive analysis of human PRP task performance. Comparison of **(A)** time series and **(B)** cumulative percent correct choices for the ET patient cohort (N=11) PRP task performance relative to simulated random task performance indicated that ET patients make optimal choices increasingly over time and significantly more often than chance overall; statistical results in (**B**) are from matched-samples t-test. (**C**) and (**D**) are same as (**A**) and (**B**) but for the behavioral cohort (N=42). There was no significant difference in cumulative percent correct choices between the ET patients and behavioral cohort (two-samples t-test, t(51) = -1.7, p = 0.09).

**Fig. S4.**
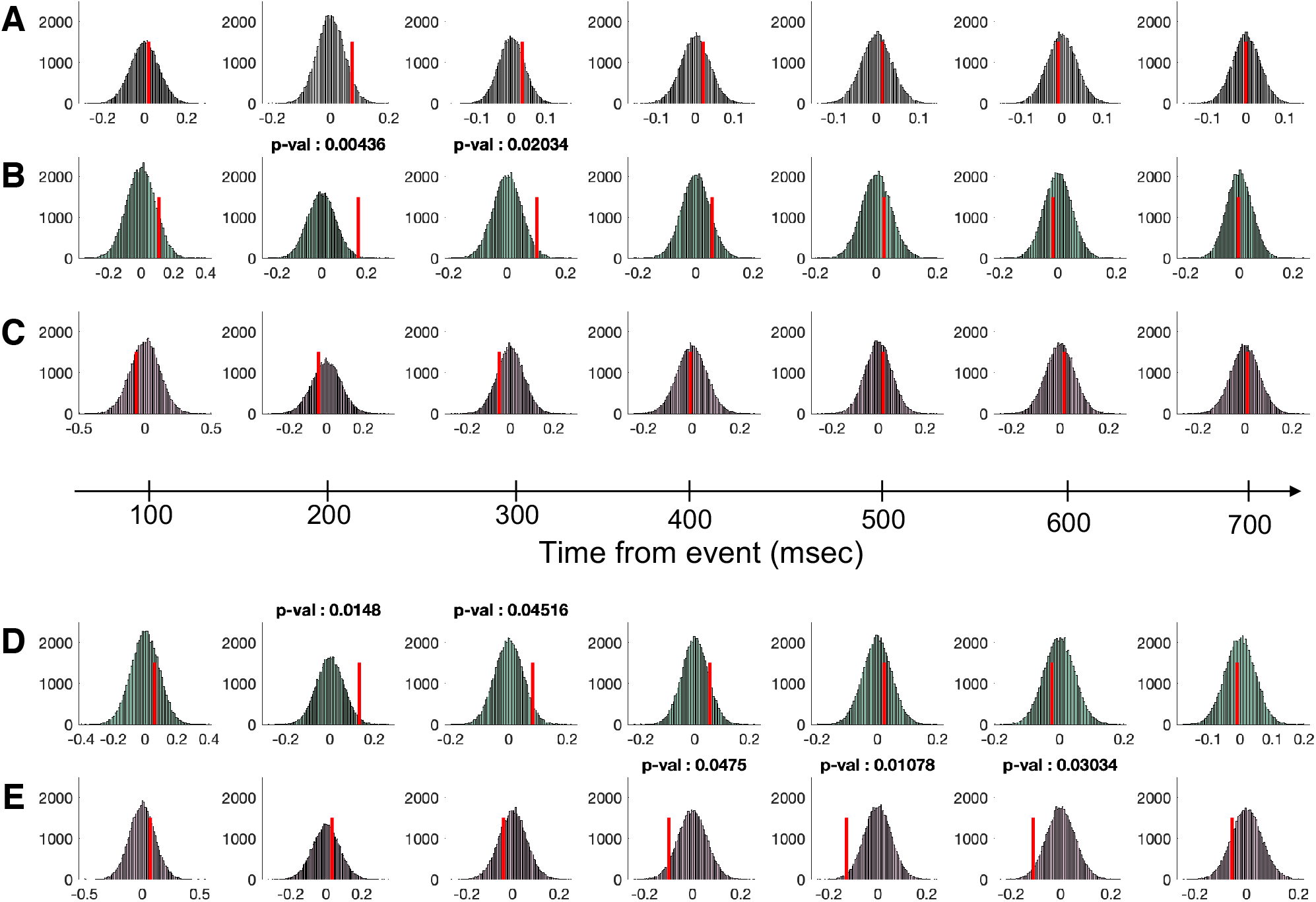
Permutation testing of differences in dopamine fluctuations in response to TDRL and VPRL prediction error signals. Comparing the output of 50,000 permutation tests (histograms) to the mean differences in the dopamine levels (vertical red bars) indicates statistically significant time points at which dopamine levels distinguish (**A**) TD-RPEs across all trials, (**B**) TD-RPEs on reward trials, (**C**) TD-RPEs on punishment trials, (**D**) VP-RPEs across all trials, and (**E**) VP-PPEs across all trials. P-values are calculated as the percentage of permutation test outputs that are greater than the actual mean difference at each time point. This non-parametric analysis recapitulated the results of the parametric analyses reported in the main text, namely that no differences were found for (**A**) TD-RPEs across all trials or for (**C**) TD-RPEs on punishment trials and that statistically significant differences were found at 200-300msec for (**B**) TD-RPEs and (**D**) VP-RPEs on reward trials and at 400-600msec for (**E**) VP-PPEs on punishment trials.

**Fig. S5.**
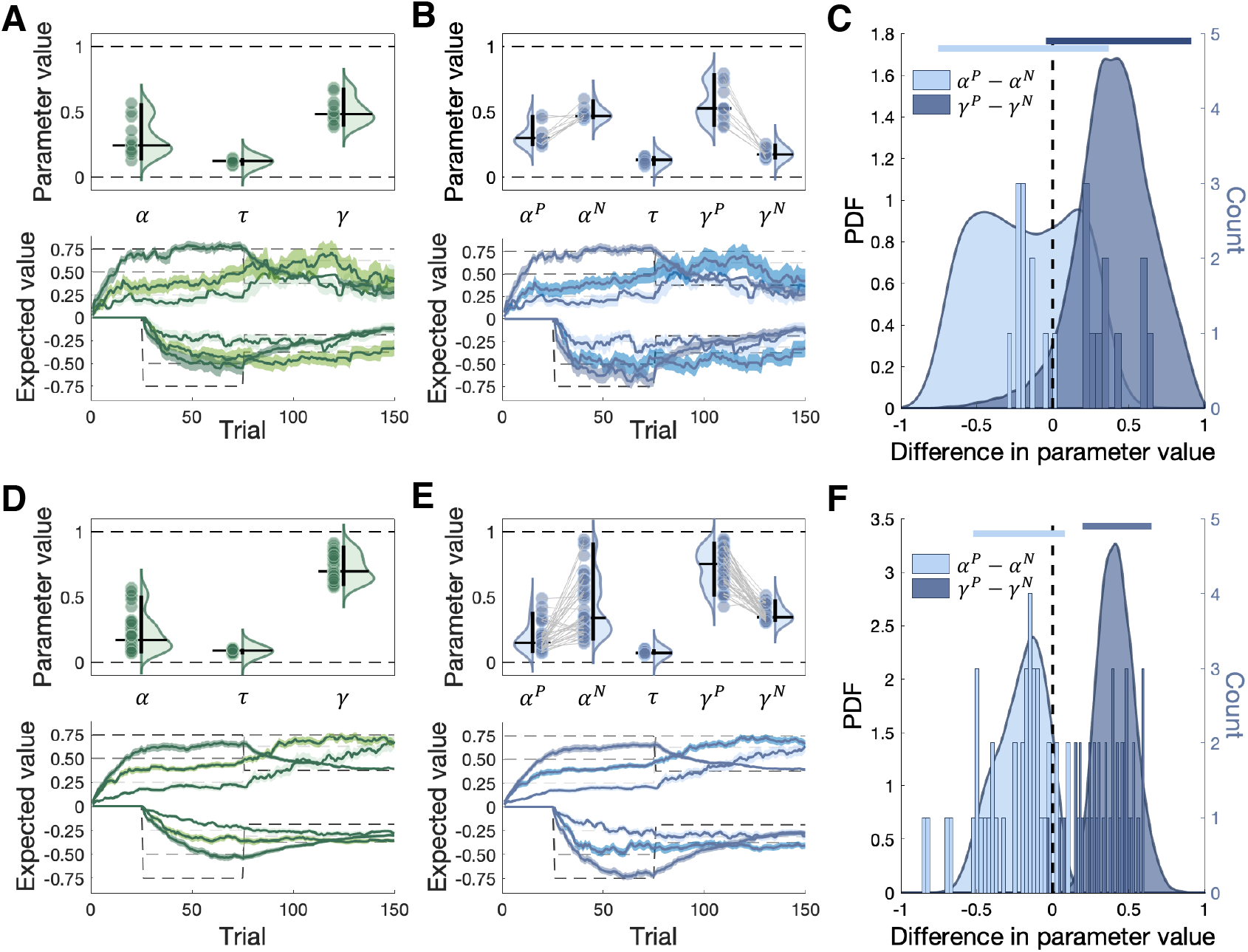
Valence-partitioning in human choice behaviors. Posterior parameter distributions for (A, top) TDRL and (B, top) VPRL models for the ET patient cohort (N=11), with individual patients’ parameter values shown as dots and the group-level distributions represented as violin plots, with the 95% highest density interval (HDI) indicated by vertical black bars. In (B, top), grey lines connect individual patient parameter values. Time series of (A, bottom) TDRL and (B, bottom) VPRL model-derived action value estimates (ribbons) plotted against the true action values (dashed grey lines); ribbons depict the mean expected value across participants (bold line) ± 1 SEM. (C) Asymmetries between reward and punishment systems in VPRL are depicted by the difference between learning rates (light blue) and temporal discount factors (dark blue) for both the group-level (distributions) and individual-level parameter values (histograms); vertical dashed black line demarcates no difference parameter distributions, horizontal blue lines depict the 95% HDI of the group-level distributions. (D)-(F) are the same as (A)-(C) but for the behavioral cohort (N=42). For the ET cohort, at the group-level (C), there were no credible differences in VPRL reward- and punishment-system parameters; at the individual-level (B), 10 out of 11 ET patients demonstrated a higher learning rate for punishments than for rewards (mean individual difference = -0.17 [-0.86 0.53]), and all 11 ET patients demonstrated a higher temporal discount factor for rewards relative to punishments (mean individual difference = 0.38 [-0.22 1.0]). For the behavioral cohort, at the group-level (F) the reward discount factor was credibly larger than the punishment discount factor 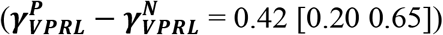 at the individual-level, 38 out of 42 participants had a larger punishment learning rate than reward learning rate (mean individual difference = -0.24 [-0.90 0.33]), and all 42 participants had a larger reward temporal discount factor than punishment temporal discount factor (mean individual difference = 0.36 [-0.19 0.85]).

**Fig. S6.**
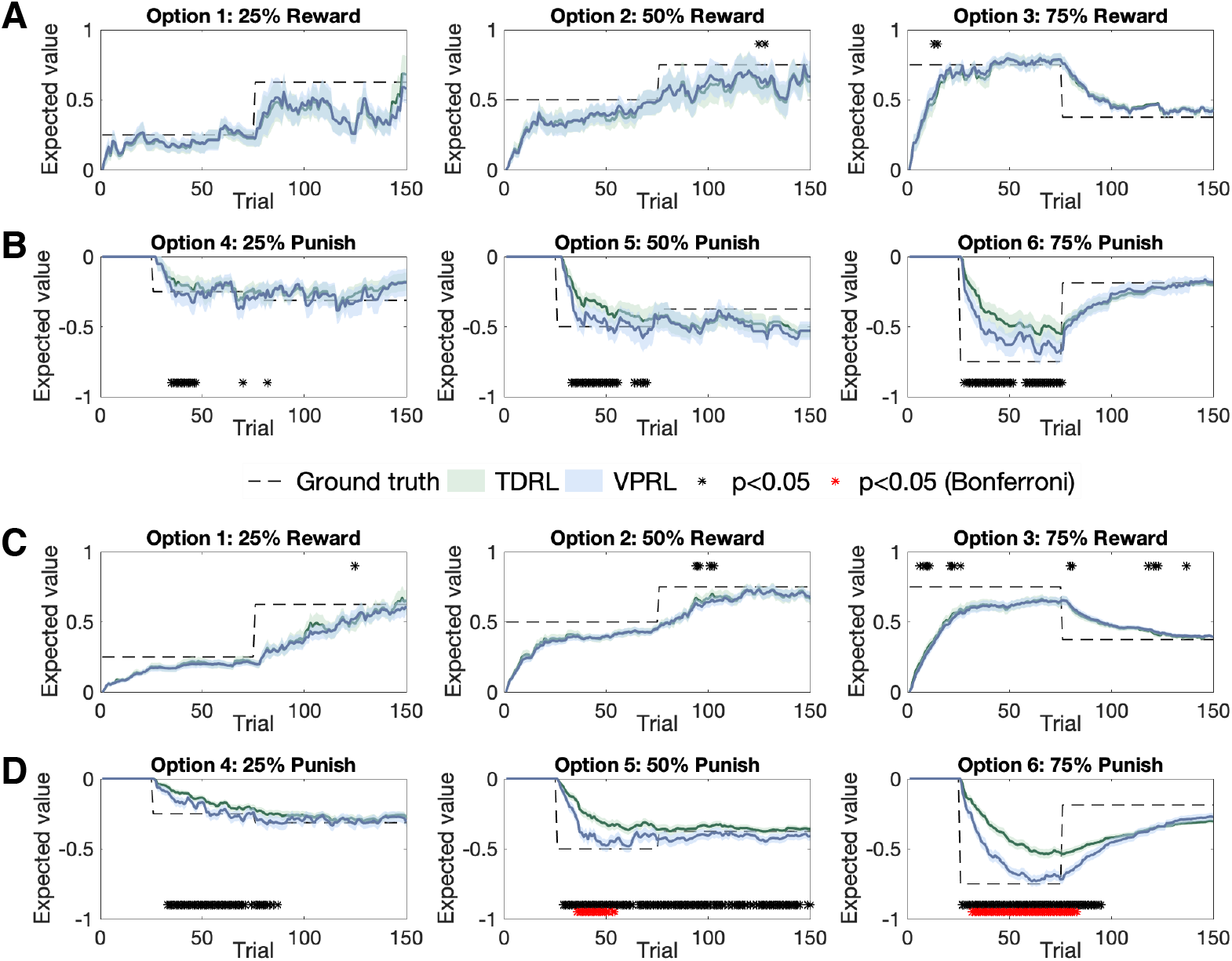
Time series of TDRL- and VPRL-derived action values for ET patients. For each participant, we used the fitted parameters for TDRL and VPRL to derive the learned action values for the rewarding and punishing options on the PRP task for (**A**,**B**) the ET cohort and (**C**,**D**) the behavior-only cohort. The bold lines in the green and blue ribbons represent the mean expected values for TDRL and VPRL across participants, respectively, and the shaded regions represent ± 1 SEM; black asterisks indicate p < 0.05 for matched-pairs t-tests, and red asterisks indicate p < 0.05 Bonferroni corrected t-tests. (**C**,**D**) are the same as (**A**,**B**) but for the behavior-only cohort. For the ET cohort, TDRL and VPRL model-derived learned values for (**A**) reward-associated options were not significantly different (two-way ANOVA (model, time): F(model) = 0.79, p = 0.38, F(time) = 2.42; p = 5.8e-14), whereas the learned values for (**B**) punishment-associated options were significantly different (two-way ANOVA (model, time): F(model) = 12.6, p = 4.3e4; F(time) = 10.6, p < 1.0e-16). Post-hoc paired-samples t-tests on the difference between the true value of each option and the TDRL or VPRL model-derived learned values across ET participants revealed that the errors of VPRL-derived learned values compared to ground-truth were significantly different from TDRL-derived learned values for punishment-associated options when they were first introduced in phase 2. For the behavior-only cohort, we found there were no significant difference between (**C**) reward-associated options (two-way ANOVA (model, time): F(model) = 3.12, p = 0.08; F(time) = 2.3, p = 6.7e-13) but did find significant differences for (**D**) punishment-associated options (two-way ANOVA (model, time): F(model) = 13.7, p = 2.0e-4); F(time) = 12.3, p < 1.0e-16), which again corresponded to differences in value estimates of punishment-associated options during phase 2.

**Fig. S7.**
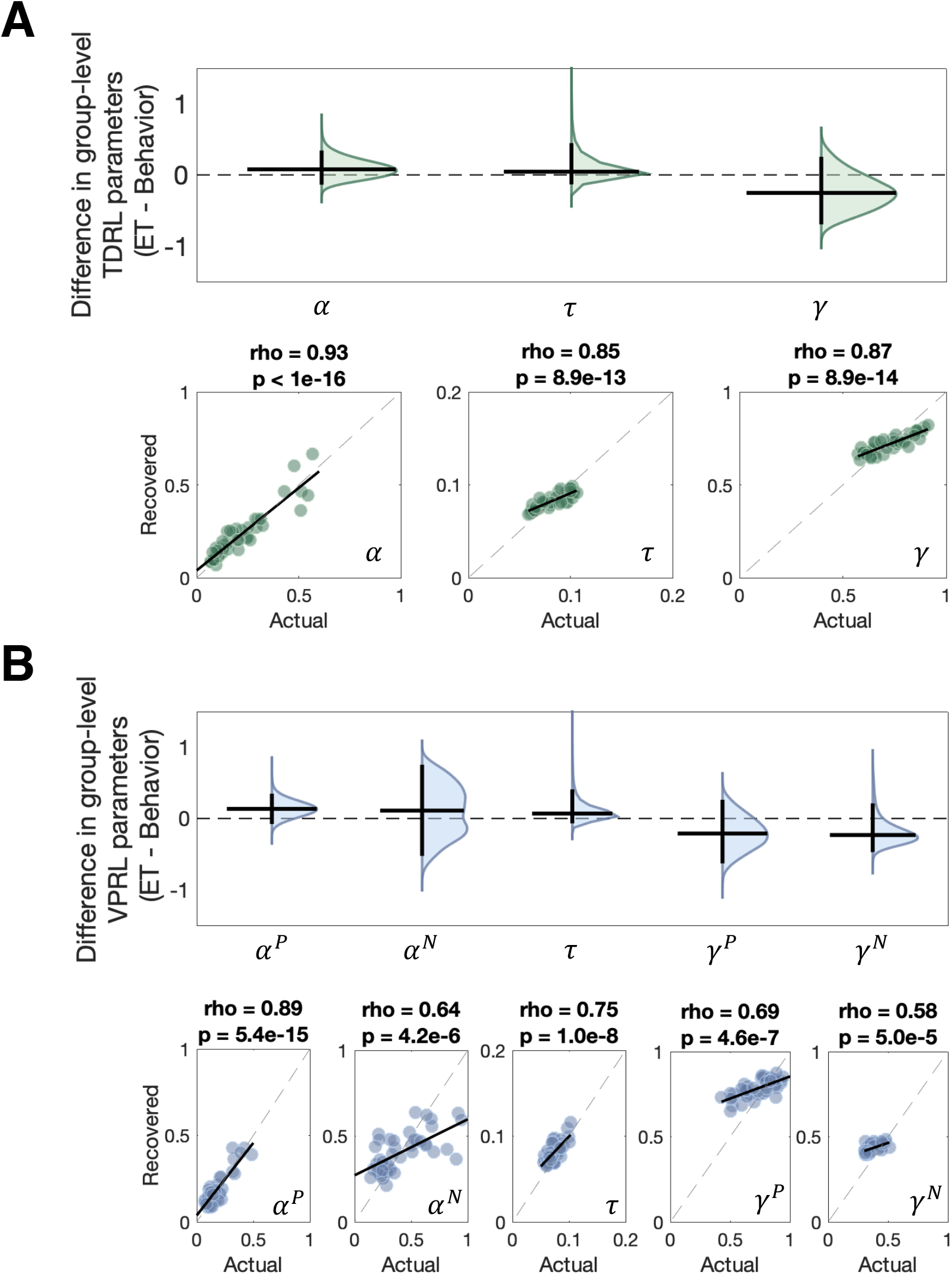
TDRL and VPRL parameter recovery. Contrasting the ET and behavioral cohort group-level parameter distributions for (**A**) the TDRL model revealed no credible differences in posterior distributions for all model parameters; horizontal black lines indicate the median value of the distribution, and the vertical black lines indicate the 95% highest density interval (HDI) of the distribution. The parameter recovery results for each TDRL model parameter is shown in the subpanel scatter plots, with rho and p-values computed using Pearson’s correlation. (**B**) is the same as (**A**) but for VPRL model parameters.

**Table S1.** Model comparison results. For both marginal likelihood (model evidence: ME) and the posterior predictive density (PD) for each model, reported values are the median estimate (log scale), with the parenthetical values reflecting either the interquartile range (model evidence) or the Monte Carlo standard error (predictive density) of estimation procedures. Note: given the hierarchical model specification, the total number of parameters for each model and each cohort is calculated as (2*K + K*N), where K is the number of unique model parameters, N is the number of participants in a cohort, the first product (2*K) represents the number of group-level parameters, and the second product (K*N) represents the number of individual-level parameters.

|  |  | Essential Tremor (n=11) |  | Behavioral (n=42) |  | Combined (n=53) |  |
| --- | --- | --- | --- | --- | --- | --- | --- |
| Model | K | ME | PD | ME | PD | ME | PD |
| TDRL | 3 | -1020.5<br>(0.09) | -1012.2<br>(31.2) | -3479.2<br>(2.7) | -3430.7<br>(95.4) | -4499.9<br>(2.0) | -4444.3<br>(105) |
| VPRL | 5 | -987.2<br>(0.09) | -975.5<br>(43.9) | -3375.4<br>(3.8) | -3319.0<br>(103) | -4113.5<br>(3.6) | -4300.1<br>(114) |
| Asymmetric TDRL | 4 | -1001.1<br>(0.1) | -988.7<br>(38.9) | -3377.4<br>(2.9) | -3326.1<br>(102) | -4177.8<br>(2.9) | -4315.7<br>(113) |
| Asymmetric VPRL | 7 | -986.3<br>(0.09) | -971.3<br>(45.0) | -3359.3<br>(2.0) | -3297.2<br>(105) | -3996.5<br>(4.0) | -4271.9<br>(117) |

**Table S2.** Model recovery results. For both marginal likelihood (model evidence; ME) and posterior predictive density (PD), reported values are the median estimate (log scale), with the parenthetical values reflecting either the interquartile range (model evidence) or the Monte Carlo standard error (predictive density) of estimation procedures. The differences in model evidence (ME), which corresponds to the Bayes Factor, and differences in predictive density (PD) are also reported as median estimates with corresponding error values; all difference measures were computed as (TDRL − VPRL), and as such positive difference values indicate greater support for TDRL and negative difference values indicate greater support for VPRL.

| True Model | Simulated Model | Essential Tremor (n=11) |  |  |  | Behavioral (n=42) |  |  |  |
| --- | --- | --- | --- | --- | --- | --- | --- | --- | --- |
| | | ME | $\Delta$ ME | PD | $\Delta$ PD | ME | $\Delta$ ME | PD | $\Delta$ PD |
| TDRL | TDRL | -933.1<br>(0.06) | 6.1<br>(0.08) | -922.3<br>(37.2) | 3.2<br>(3.6) | -3214.9<br>(0.2) | 31.3<br>(0.35) | -3175.2<br>(86.6) | 28.2<br>(6.2) |
|  | VPRL | -939.2<br>(0.06) |  | -925.5<br>(36.5) |  | -3246.2<br>(0.3) |  | -3203.4<br>(85.5) |  |
| VPRL | TDRL | -962.4<br>(0.07) | -9.6<br>(0.12) | -952.5<br>(38.6) | -13.8<br>(9.5) | -3166.8<br>(0.27) | -36.9<br>(0.46) | -3137.0<br>(84.9) | -47.0<br>(14.6) |
|  | VPRL | -952.8<br>(0.08) |  | -938.7<br>(42.0) |  | -3129.8<br>(0.36) |  | -3090.0<br>(87.5) |  |

## Notes

### Competing Interest Statement

The authors have declared no competing interest.

